# Anticipatory regulation of cardiovascular system on emergence of auditory-motor interaction in young infants

**DOI:** 10.1101/2021.10.11.464010

**Authors:** Yuta Shinya, Kensuke Oku, Hama Watanabe, Gentaro Taga, Shinya Fujii

## Abstract

Humans develop auditory-motor interaction to produce a variety of rhythmic sounds using body movements, which are often produced and amplified with tools, such as drumming. The extended production of sounds allows us to express a wide range of emotions, accompanied by physiological changes. According to previous studies, even young infants enhance movements in response to auditory feedback. However, their exhibition of physiological adaptation on emergence of auditory-motor interaction is unclear. We investigated the movement and cardiac changes associated with auditory feedback to spontaneous limb movements in 3-month-old infants. The results showed that infants increased the frequency of limb movements inducing auditory feedback, while they exhibited the more regular rhythm of the limb movements. Furthermore, heart rate increase associated with the limb movement was first inhibited immediately after the timing of the auditory feedback, which may reflect sustained attention to the auditory stimuli. Then, through auditory-motor experience, the heart rate increase was inhibited even prior to the auditory feedback, leading to suppression on the peak intensity of the heart rate increase. These findings suggest that young infants regulate the cardiovascular system as well as limb movements in anticipation of the auditory feedback. The anticipatory regulation associated with movement and attentional changes may contribute to reduced cardiovascular stress in auditory-motor interaction, and provide a developmental basis for more sophisticated goal-directed behavior of producing rhythmic sounds.

## Introduction

Humans produce a variety of sounds not only by using their vocal organs, but also their body movements, such as through drumming. The beat is often produced and amplified with tools such as musical instruments (Patel 2014); for example, tool-assisted rhythmic drumming is assumed to be unique to humans, except for only a few species of animals (e.g. palm cockatoo) (Heinsohn et al. 2017). The extended production of sounds allows us to express various rhythms and melodies to induce a wide range of emotions and feelings, accompanied by physiological as well as behavioral changes (Blood and Zatorre 2001; Habibi and Damasio 2014; Scherer and Zentner 2001). Furthermore, in the long term, these changes associated with music activities are assumed to play a crucial role in subjective well-being and health through homeostatic processes, such as stress reduction (Bainbridge et al. 2020; Chanda and Levitin 2013; Cirelli and Trehub 2020; de Witte et al. 2020; Tramo et al. 2011).

Recent cognitive neuroscientific research of music shows that both perception and production of music require the interaction between auditory and motor systems (Zatorre et al. 2007). Especially, the integration of spatial, auditory, and motor information has been assumed to be essential in producing rhythmic sounds by controlling movements, such as drumming. Humans acquire universal musicality, including the ability to produce rhythmic sounds, which is often amplified using musical instruments (Kotz et al. 2018; Savage et al. 2015). However, the emergence of these auditory-motor interactions in the acquisition of sound production in early development remains unclear.

As opposed to sound perception, which indicates the early ability to process musical sounds (e.g. Hannon and Johnson 2005; Perani et al. 2010; Phillips-Silver and Trainor, 2005; Winkler et al. 2009), the development of sound production in infancy is directly revealed by relatively few studies. From early developmental stages, human infants produce rhythmic movements, typically defined as repeated body movements in the same form at regular short intervals (e.g. banging, shaking, ringing) (Ejiri 1998; Thelen 1979; Zentner and Eerola 2010), and more recently, they have also been reported to exhibit a certain degree of tempo flexibility (Cirelli and Trehub 2019; Fujii et al. 2014; Provasi et al. 2014; Rocha and Mareschal, 2017; Yu and Myowa, 2021; Zentner and Eerola 2010). For example, Zentner and Eerola (2010) reported that 5-to 24-month-old infants exhibited more rhythmic movements to musical or rhythmic stimuli than to speech, and they moved faster when the tempi were faster. Rocha and Mareschal (2017) also reported that infants at 18-months-old modulated bell ringing to the music tempo, although they showed no evidence of synchronous movements. Fujii et al. (2014) reported that three-to four-months-old infants are already primed to interact with music via limb movements and vocalizations. Therefore, it is possible that even young infants develop the precursors of auditory-motor interactions, allowing for more sophisticated music production such as dancing or drumming in later developmental stages.

Previous studies investigating the development of memory or sense of self-agency have shown that young infants would enhance movements in response to contingent changes in the environment including auditory sounds (Bahrick and Watson 1985; DeCasper and Fifer 1980; Lee and Newell 2013; Rochat and Striano 1999; Rovee-Collier 1999; Watanabe et al. 2011). For example, in a situation where sucking induced presentation of mother’s speech or face, new-borns increased their oral movements to induce the presentation (DeCasper and Fifer 1980). Similarly, in the sucking task, 2-month-old infants showed modulation of their oral movements in conditions where the pitch of the contingent auditory feedback varied depending on the pressure of their sucking (Rochat and Striano 1999). Furthermore, in the mobile task, supine infants at 3-months-old with a limb connected to an overhead mobile selectively increased the limb’s movements to induce audio-visual stimulation from the mobile (Watanabe et al. 2011).

These findings suggest that young infants are motivated to increase their body movements to produce sounds in situations where their body movements could induce or be converted to auditory information. Noteworthily, in this period, it would be challenging for infants to inhibit and regulate spontaneous movements automatically generated by subcortical systems (i.e. central pattern generator in brainstem and spinal cord) (Watanabe et al. 2011; Prechtl 1997). Thus, the findings are assumed to be evidence for the precursor of goal-directed behavior (e.g. reaching; Lee and Newell 2013) due to cortical development, providing a base for the acquisition of music production.

As opposed to behavioral adaptation (i.e. enhancement of motor activities), few studies investigated the physiological activities through the auditory-motor interaction in early infancy. Reportedly, foetus (Lecanuet et al. 1992) and young infants (Morrongiello and Clifton 1984; Porges et al. 1973) display heart rate deceleration when exposed to auditory stimuli, which may be a function of orienting response (Graham and Clifton 1966) and sustained attention (Richards and Casey 1991). Also, such cardiovascular response may depend on predictability of the timing of auditory stimuli (Hajcak et al. 2003; 2004). Given that the auditory-motor interaction allows prediction of auditory feedback in response to movement (Zatorre et al. 2007), it is possible that the dynamic change in cardiovascular system may occur in the early process of auditory-motor interaction in young infants.

Moreover, as a notable aspect of music, its production and perception elicit feelings and emotions which involve arousal, stress regulation, and behavioral and attentional changes (Habibi and Damasio 2014; Scherer and Zentner 2001). Especially, auditory-motor synchronization has been reported to decrease the physiological load in cardiovascular system, even in the absence of musical component (Bood et al. 2013; Chaisurin et al. 2020; Trappe 2010). This predictive mode to maintain homeostasis regulation via behavioral and physiological changes is considered as ‘allostasis’ (Sterling 2012). Therefore, we speculate that the auditory-motor interaction in young infants would accompany not only behavioral adaptation, but also dynamic change in the cardiovascular system.

To the best of our knowledge, no study has revealed the dynamic change in cardiovascular activity in response to auditory feedback associated with movement in infants. A few studies investigated more global changes of parasympathetic activities, indexed by respiratory sinus arrhythmia (RSA; i.e. high frequency of heart rate variability [HRV]) (Task Force 1996; Denver et al. 2007), in the mobile paradigm using audio-visual feedback (Haley et al. 2008; Sullivan 2016). Haley et al. (2008), reported that, 3-month-old infant learners showed greater suppression of RSA than the non-learners. Alternatively, Sullivan (2016) reported that, in 5-month-old infants, RSA did not vary significantly over sessions, and was not associated with learning status. Nevertheless, more localized, and dynamic cardiovascular changes associated with auditory-motor experience in young infants remains unclear.

By focusing on the physiological aspects, the current study expands the previous observation on emergence of auditory-motor interaction in early infancy (Bahrick and Watson 1985; DeCasper and Fifer 1980; Rochat and Striano 1999; Rovee-Collier 1999; Watanabe et al. 2011). To reveal the physiological adaptation in this context, we investigated cardiac as well as movement changes through auditory feedback to limb movements in 3-month-old infants. Considering the role of sound production and perception in achieving homeostasis via arousal and stress regulation (Bainbridge et al. 2020; Chanda and Levitin 2013; Cirelli and Trehub, 2020; de Witte et al. 2020; Tramo et al. 2011; Sterling 2012), we hypothesized that the heart rate change related to limb movements inducing auditory feedback may be regulated during auditory-motor integration progress, as well as changes in limb movements.

## Methods

### Participants

Thirty 3-month-old term infants (10 females and 20 males) participated in this study. The sample excluded preterm/low-birth weight infants or infants with medical problems. The mean age of the infants was 99.77 days (range: 90–110 days, SD = 5.54 days). Twenty-seven other infants participated but did not complete the experiment due to excessive crying (*n* = 25) and drowsiness (*n* = 2). An additional 10 infants completed the experiment, but were excluded because their excessive fussiness was soothed by the experimenter occasionally during the period of experiencing auditory feedback (i.e., Test Phase; see Experimental procedure section). The study was approved by the ethical committee of Life Science Research Ethics and Safety, The University of Tokyo, and was conducted in accordance with standards specified in the 1964 Declaration of Helsinki. Written informed consent was obtained from the participants’ parents.

### Apparatus and stimuli

Infants were studied in the laboratory at the Department of Physical and Health Education at the Graduate School of Education, the University of Tokyo. During the experiment each infant was positioned in a supine position on a baby mattress (120 cm × 70 cm).

Four virtual drum-kit sensors with accelerometer and gyroscope (Freedrum, Sweden [c.f. https://freedrum.rocks]; Fig. 1*a*) were put on each limb of the participants to record the timing of the acceleration of each limb’s movement exceeding a threshold. When the threshold was exceeded during Test phase, auditory stimuli was immediately fed to an infant from two speakers (Bose) about 40 cm near their ears. The auditory stimuli produced by each limb’s sensor corresponded to drum sounds (i.e., high Tom, mid Tom, low Tom, floor Tom; Fig. 1*b*) with a duration of 2 sec and varying in heights [pitch = 178.3, 143.5, 114.2, 92.86 Hz, respectively] of approximately 60 dB. These stimuli were obtained from the instrument source in music production software (the category ‘Session Dry Kit’ in ‘Ableton live 9 suit [Ableton]’).

**Fig. 1.**
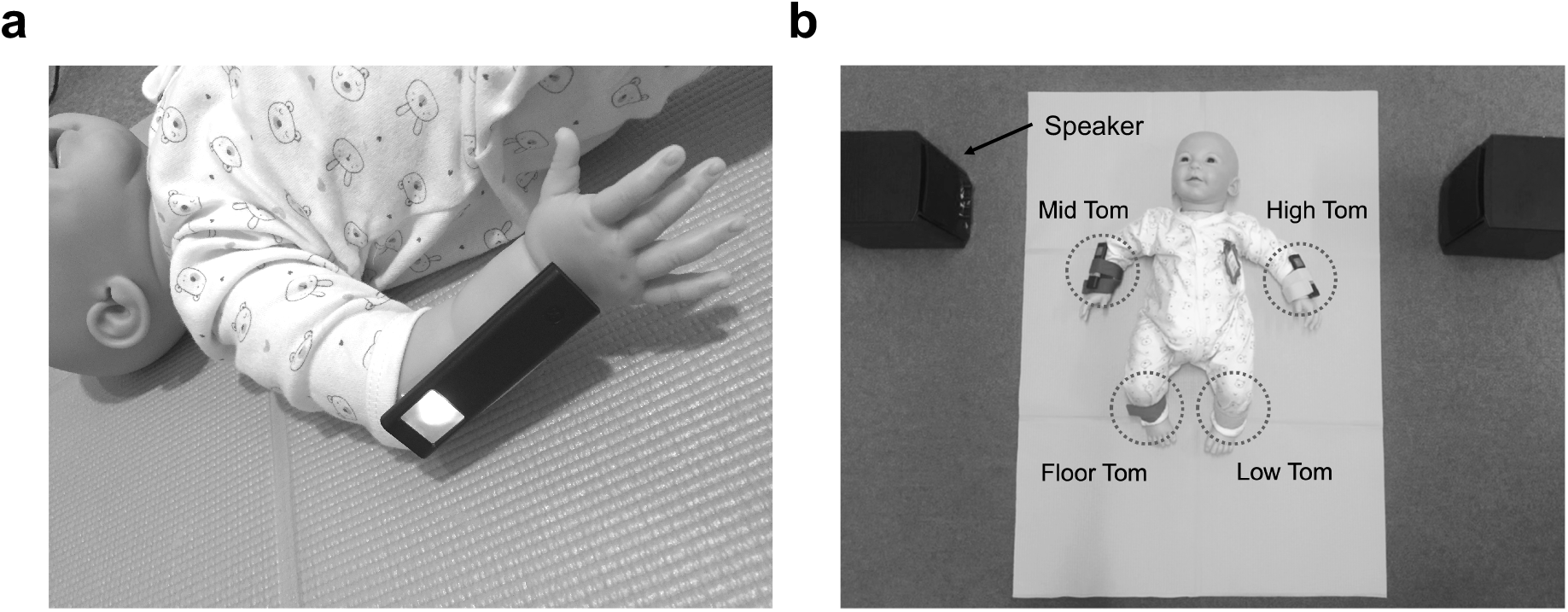
Experimental setup: an infant wearing a virtual drum kit sensor (Freedrum, Sweden) on the right arm (**a**); the sounds produced by each limb’s sensor correspond to drum sounds of different heights (**b**).

For heart rate recording, we used a wireless multi-telemeter (WEB-1000, Nihon Koden) to obtain the Electrocardiograph (ECG) data. The ECG data were measured from three disposable paediatric ECG electrodes (NC Vitrode, Nihon Koden) using lead II (right collarbone and lower left rib), grounded at the left collarbone. The sampling rate was 1000 Hz.

Additionally, a single video camera (Handycam HDR-CX675, SONY) was used to record the movements, faces, voices, and status of the sensors to assess infants’ behavioral status and maintain records.

### Experimental procedure

Infants were positioned on their backs on the baby mattress before commencing the experiment. Four virtual drum-kit sensors were attached to the forearms and the lower legs of each infant.

The experimental session consisted of three periods lasting for a total of 12-min: Pre phase (2-min), Test phase (8-min), and Post phase (2-min). During Pre and Post phases, the speakers’ volume was muted to not feed the auditory stimulus contingent to the spontaneous limb movements to the infants. During Test phase, infants were able to listen to and induce the auditory feedback through limb movements. For statistical analysis, Test phase was further divided into four two-minute sub-phases (T1, T2, T3, T4).

During this procedure, the caregiver stayed away from the infant’s view and the experimenter did not interact with the infant socially (e.g. eye-contact, speech) to avoid distracting the infant. When the infant became excessively fussy or upset, the experimenter attempted to soothe the infant, in line with previous studies (e.g., Haley et al. 2008). However, before Post phase, the infant was excluded from the analysis for excessive crying because their emotional state could be due to factors other than auditory feedback (e.g., sleepiness, fatigue, fear of strangers, etc.) and would affect movement and cardiac measures. On the other hand, during Post phase (i.e. when the auditory feedback disappeared), a relatively high number of infants showed signs of fussiness or upset, so these infants were included in the analysis.

### Data analysis

#### Movement measures

We obtained the onset time of all limb movements which exceeded the threshold to induce auditory feedback from all participants (*n* = 7400). If the time difference from the previous data within the same limb was less than 100 msec, that data was excluded as a double count from analysis (*n* = 318). Then, if the timing was the same among different limbs, it was considered a single count, and the same timing data was excluded (*n* = 225). We also detected the period when the sensors had come off an infant’s limb using video data, and the timing data related to them were excluded (*n* = 121). Thus, we used a total of 6736 timing data of the movement for calculating movement measures. Based on the time series of the timing data, we calculated the frequency of the limb movements and the coefficient of variation (CV) of the time intervals between the limb movements for 2-min for each infant to assess the changes in tempo and regularity of limb movements (Rocha et al. 2021).

#### Cardiac measures

The recorded ECG data were converted into R-wave intervals after manual artifact correction with Heart Rate Variability Analysis Software (Mindware Technologies LTD, US). The corrected R-wave intervals were converted into time series of heart rate.

Power spectral analysis of heart rate variability (HRV) was performed using the Kubios HRV Analysis Software 2.0 (The Biomedical Signal and Medical Imaging Analysis Group, Department of Applied Physics, University of Kuopio, Finland) (Tarvainen et al. 2014). We determined two main oscillations (Task Force 1996; Denver et al. 2007): a low-frequency component (LF, 0.04–0.24Hz), representing both sympathetic and parasympathetic activity related to the baroreflex system; and a high-frequency component (HF, 0.25–1.50 Hz) representing vagal (parasympathetic) activity modulated by respiratory cycles (i.e., RSA). For the high-frequency component, the bandwidth was extended to 0.25–1.50 Hz from the adult standard of 0.15–0.40 Hz, due to the higher speed of respiration in young infants (Shinya et al. 2016). Both HRV components were calculated by summing power spectral density values over the bandwidth. The ratio of LF to HF was also calculated as an indicator of sympathovagal balance.

In addition to directly revealing the immediate heart rate change due to the contingent auditory feedback of limb movements, we determined the event-related heart rate change (ERHR) by extracting heart rate change for 10 sec before and after each timing for each limbs’ acceleration exceeding the threshold (i.e. the onset of auditory feedback during Test phase). The ERHR baseline was adjusted by subtracting the cubic trend of the time series of heart rate for each phase and each participant. In this analysis, we used a total of 6679 timing data after excluding the timing data of less than 10 seconds after the experiment start (*n* = 57) from the timing data used in the analysis of the movement measures.

### Statistical analysis

To assess the global changes of the movement and cardiac measures except for ERHR, we conducted a within subjects one-way analysis of variance (ANOVA, phase: Pre, T1, T2, T3, T4, Post). In the post-hoc analysis, we examined the effects of each phase from Pre phase to Test and Post phases, and used sequentially rejective Bonferroni’s corrections for the multiple comparisons.

To assess the statistically significant occurrence of ERHR for each phase, we created surrogate data by randomly extracting data 200 times from the timeseries of each infant’s heart rate. For each phase (i.e., Pre, T1, T2, T3, T4, and Post) of the waveforms of ERHR (original: *n* = 814, 716, 867, 1080, 1334, 1868, respectively [total *n* = 6679]; surrogate: each *n* = 6000), we performed linear mixed-effects models (LMM) including the fixed effect of event (i.e., original vs surrogate) and the random effect of participant (*n* = 30) with intercept to test whether significant difference occurred at each 0.1 sec time point between the original and surrogate ERHRs. As the LMMs were repeated a total of 201 times in each phase, sequentially rejective Bonferroni procedures were applied for the multiple testing corrections.

We also calculated difference-waveforms of ERHR in T1–T4 and Post phases by subtracting the averaged waveforms of ERHR in Pre phase for each participant to examine the significant change through auditory-feedback. We performed the same LMMs as the above analysis for each 0.1 sec time point in each phase (i.e. T1, T2, T3, T4, and Post) of the difference-waveforms of ERHR (original: *n* = 716, 867, 1080, 1334, 1868 [total *n* =5865], respectively; surrogate: each *n* = 6000).

Additionally, to investigate how the amplitude of ERHRs changes across phases, we conducted the LMMs to predict the mean ERHR for the period 0-2 sec after each timing for each limbs’ acceleration exceeding the threshold (*n* = 6679) by phase (i.e., Pre, T1, T2, T3, T4, post) as a fixed effect and participant with intercept (*n* = 30) as a random effect. In the post-hoc analysis, we compared the mean ERHR between Pre phase and the other phases, and sequentially rejective Bonferroni’s corrections were performed for the multiple comparisons.

The LMMs allowed us to assess the occurrence and change of ERHR across phases without assuming that the observations for successive events were independent as a general linear model (e.g., ANOVA) would require (Quené and van den Bergh 2004). Also, the models allowed for significance tests with high statistical power, as the event-related response data were not averaged over phases per participant.

All statistical analyses were performed using R version 4.0.1 (R Development Core Team 2020). The LMMs were developed using the ‘lme4’ package (Bates et al. 2015).

## Results

### Change of movement and cardiac measures across phases

Fig. 2 shows the changes of the averaged movement (Fig. 2a) and cardiac measures (Fig. 2b) across 6 phases (i.e., Pre, T1, T2, T3, T4, Post phases).

**Fig. 2.**
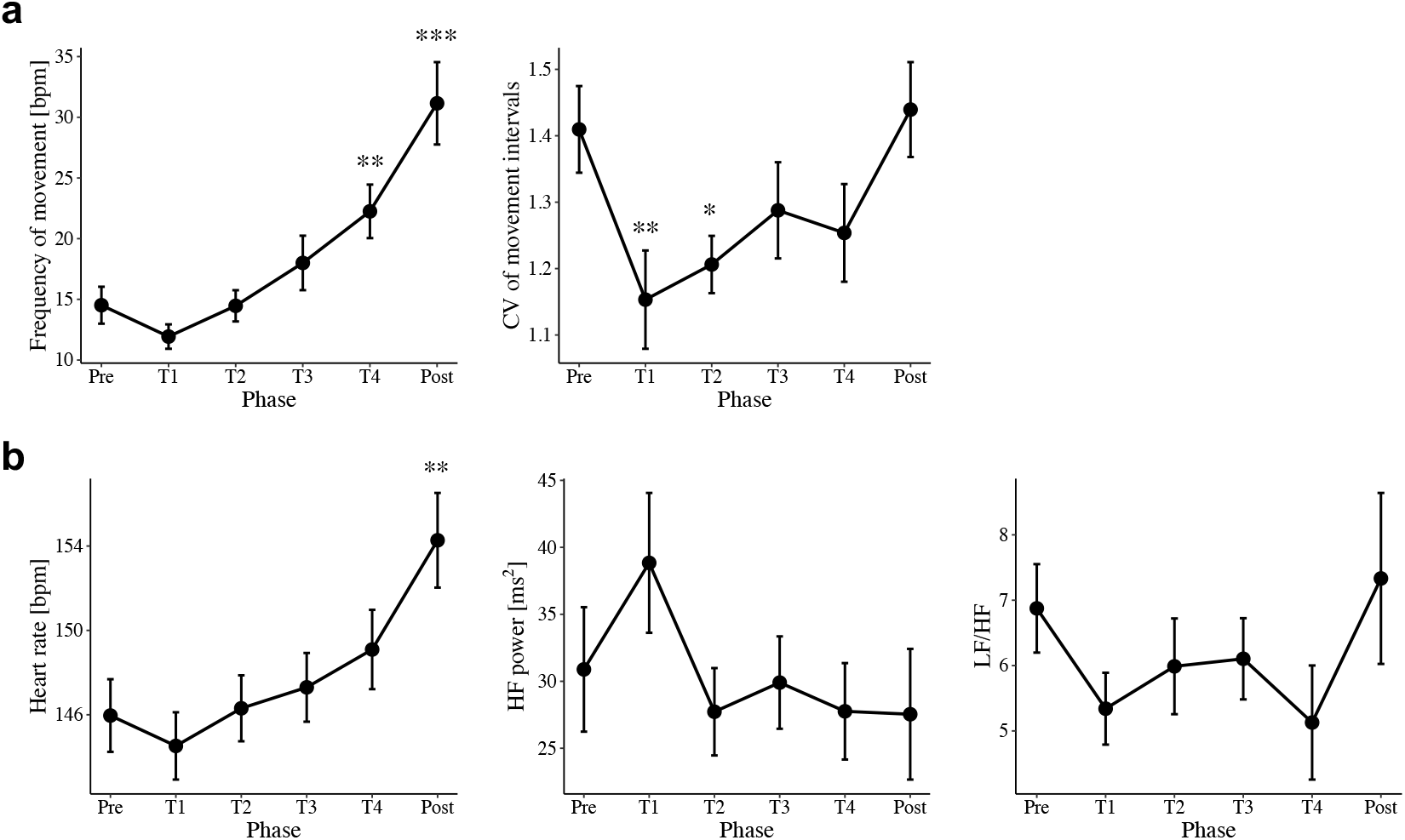
Line plots for changes of (**a**) movement measures (Frequency of movement, CV of movement intervals) and (**b**) cardiac measures (heart rate, LF/HF, HF) across phases. Bars represent standard errors. ^***^*p* < .001, ^**^*p* < .01, ^*^*p* < .05 (compared to Pre phase).

For the movement frequency related to auditory feedback, we found a significant main effect of phase (*F*_*5,145*_ = 18.31, *p* < .001). Multiple comparisons of phases revealed that the frequency of the movements during T4 and Post phases was greater than those during Pre phase (*p* = .003; *p* < .001). On the other hand, for the CV of the movement intervals, we found a significant main effect of phase (*F*_*5,145*_ = 3.388, *p* = .006). The CV of the movement intervals during T1 and T2 phases was lower than those during Pre phase (*p* = .007; *p* = .032). These results indicate that infants increased the frequency of limb movements and decreased the variance of limb movement intervals during the auditory feedback.

We observed a significant main effect of phase for heart rate (HR) (*F*_*5,145*_ = 13.24, *p* < .001). In multiple comparisons, the HR during Post phase was greater than those during Pre phase (*p* = .003). For HRV measures, only a significant main effect of phase on HF was observed (*F*_*5,145*_ = 2.48, *p* = .034), but there was no significant difference in multiple comparisons. These results indicate that, although infants showed increased HR during Post phase, the relatively stable heart rate’s and sympathetic/parasympathetic activity’s changes were observed from Pre to Test phases.

### Comparisons of ERHR waveforms between original and surrogate data

Fig. 3a shows the averaged original and surrogate waveforms of the event-related heart rate change (ERHR) in Pre, T1–T4, and Post phases for all participants. To determine the time point of significant ERHR occurrence, we performed the LMMs including the fixed effect of event (i.e. original vs surrogate) and the random effect of participant with intercept for each 0.1 second of ERHR.

**Fig. 3.**
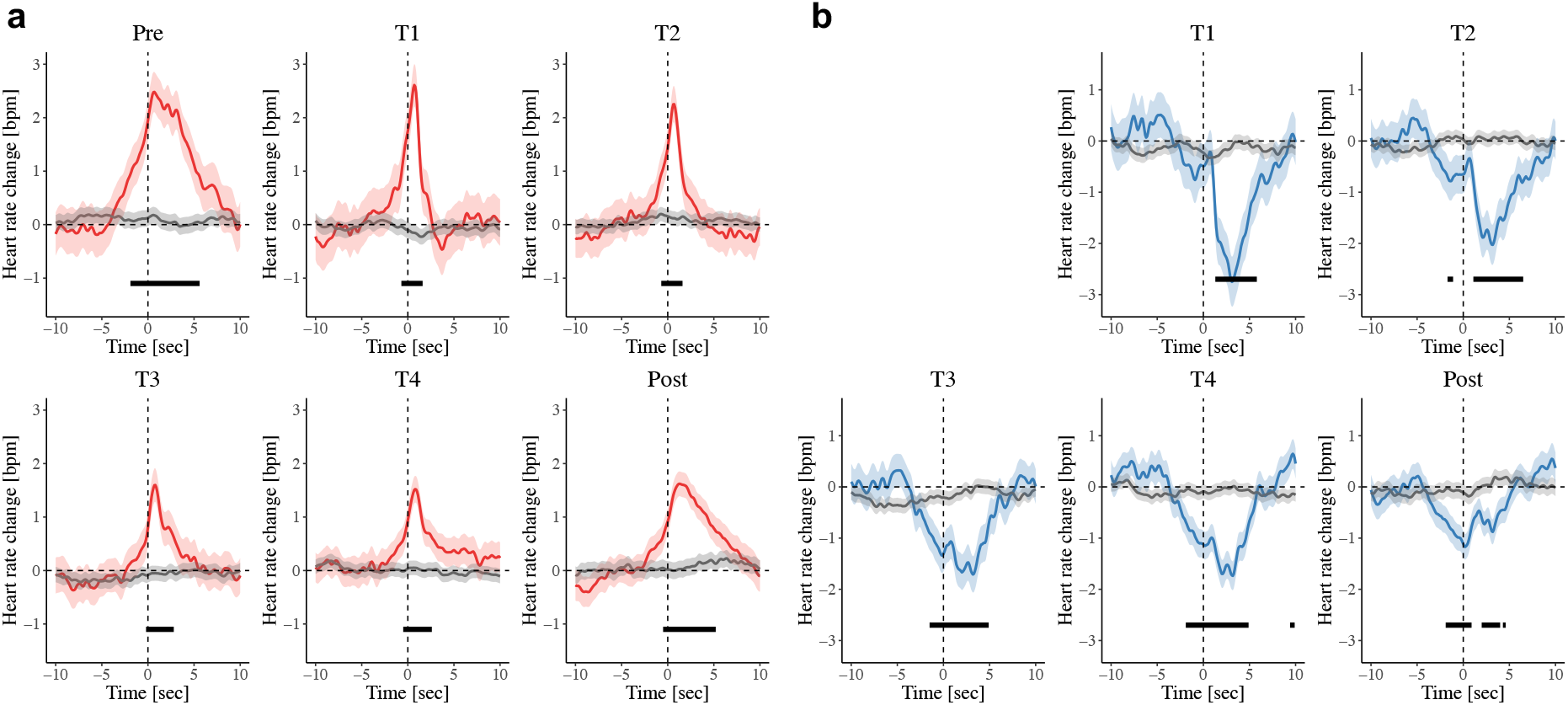
**a** Average waveforms of event-related heart rate change (ERHR) for each limb’s acceleration exceeding the threshold (i.e. onset of auditory feedback during Test phase) for each phase (Pre, T1, T2, T3, T4, Post; original [red]: *n* = 814, 716, 867, 1080, 1334, 1868, respectively; surrogate [grey]: each *n* = 6000). **b** Average difference-waveforms of ERHR between Pre and each phase (T1, T2, T3, T4, Post phase; original [blue]: *n* = 716, 867, 1080, 1334, 1868, respectively; surrogate [grey]: each *n* = 6000). Shadows represent 95% confidence intervals, and black bars represent significant differences between original and surrogate data (adjusted *p* < .05).

The models revealed that the original ERHR was significantly larger than surrogate one during -1.7 ∼ 5.1 sec in Pre phase. Alternatively, in Test phase, the periods of significant difference were relatively shorter compared to Pre phase (T1: -0.7 ∼ 1.6 sec; T2: -0.7 ∼ 1.6 sec; T3: -0.2 ∼ 2.8 sec; T4: -0.5 ∼ 2.6 sec). The start point of the significant increase in heart rate in Post phase was closer to the onset, although the end point was similar than that in Pre phase (−0.6 ∼ 5.3 sec). These results indicate that experiencing auditory feedback on limb movements led to increase and decrease in heart rate closer to the onset.

### Difference-waveforms of ERHR between Pre phase and each phase

Fig. 3b shows the averaged difference-waveforms of the ERHR in T1-T4 and Post phase, by subtracting the averaged ERHR in Pre phase. To investigate the time point of significant ERHR change from Pre phase, we performed the LMMs including the fixed effect of event (i.e. original vs surrogate) and the random effect of participant with intercept for each 0.1 second of ERHR.

We found that, in T1 phase, the original ERHR was significantly lower than the surrogate ERHR immediately after the onset (1.3 ∼ 5.8 sec), indicating that infants showed decreased heart rate in this period compared to Pre phase. After T1 phase, heart rate decreased significantly both before and after the onset (T2: -1.7 ∼ -1.1, 1.1 ∼ 6.5 sec; T3: -1.5 ∼ 4.9 sec; T4: -1.9 ∼ 4.9 sec; Post: -1.0 ∼ 0.9, 2.0 ∼ 4.0, 4.3 ∼ 4.6 sec). These results indicate that by experiencing the auditory feedback, infants first decreased heart rate after the onset, and then inhibited heart rate before the onset.

### Change in ERHR’s peak intensity

Fig. 4 shows the change in ERHR’s peak intensity (0–2 sec period) across phases. The LMM revealed a significant phase effect (*F*_*5,6671*.*8*_ = 6.14, *p* < .001). Multiple comparisons indicated that ERHR’s peak intensity during Test phase was significantly lower than that during Pre phase (T1: *t* = 2.88, *p* = .044; T2: *t* = 3.40, *p* = .008; T3: *t* = 4.78, *p* < .001; T4: *t* = 4.95, *p* < .001). Moreover, the peak intensity during Post phase was significantly lower than Pre phase (*t* = 4.45, *p* < .001).

**Fig. 4.**
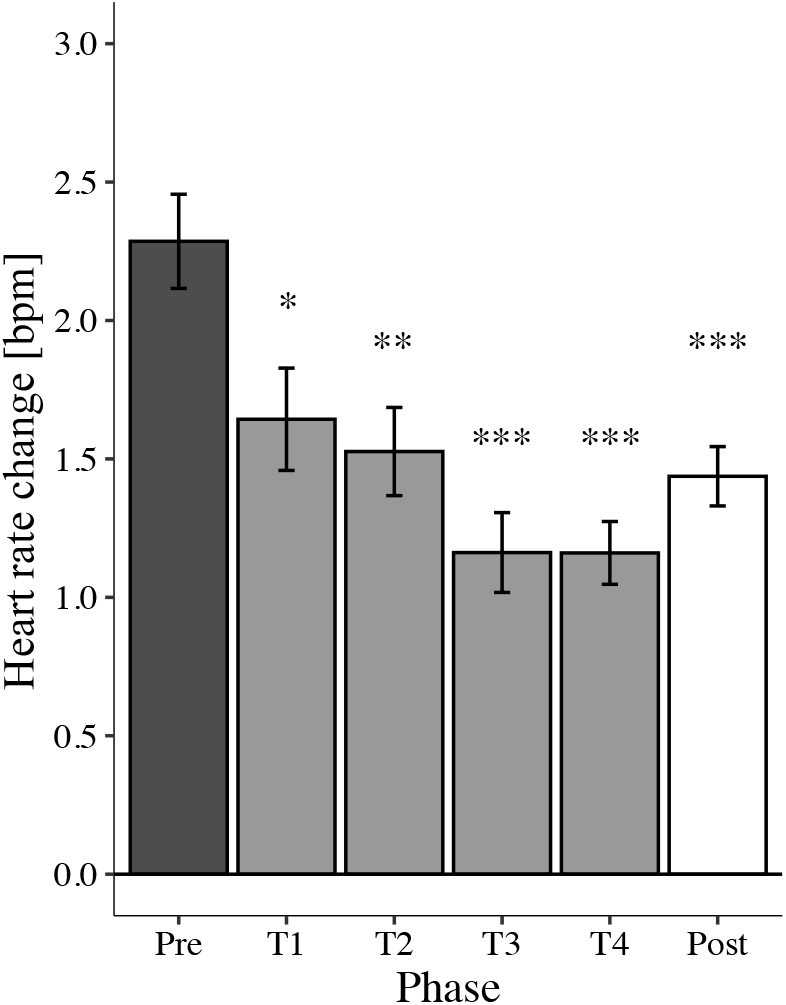
Bar plots of event-related heart rate change’s peak intensity (0–2 sec period) across Pre (black), Test (T1–T4; grey), and Post phase (white). Bars represent standard errors. ^***^*p* < .001, ^**^*p* < .01, ^*^*p* < .05 (compared to Pre phase).

## Discussion

The current study is the first to reveal how cardiovascular system would be dynamically regulated on emergence of auditory-motor interaction in early infancy, expanding previous observations regarding behavioral adaptation (Bahrick and Watson 1985; DeCasper and Fifer 1980; Rochat and Striano 1999; Rovee-Collier 1999; Watanabe et al. 2011). We found that, in 3-month-old infants experiencing auditory feedback to spontaneous limb movements, they inhibited the heart rate increase associated with the limb movements even before the auditory feedback, leading to the suppressed peak intensity of the heart rate increase. This new evidence on physiological adaptation suggests that young infants regulate the cardiovascular system as well as limb movements in anticipation of the auditory feedback.

Several studies investigating memory or sense of self-agency in young infants have reported behavioral adaptation to environmental changes including auditory feedback (i.e. enhancement of motor activities) (Bahrick and Watson 1985; DeCasper and Fifer 1980; Rochat and Striano 1999; Rovee-Collier 1999; Watanabe et al. 2011). Consistent with these findings, we confirmed that 3-month-old infants significantly increased frequency of limb movements inducing auditory feedback from Pre phase to Test and Post phases. On the other hand, little is known about how limb movement rhythms would change by experiencing the contingent feedback, although the tempo itself increases. We found that infants exhibited a significant decrease in variance of limb movement intervals during Test phase (especially, during T1 and T2), suggesting that the limb movements changed from more disorganized to more regular during the experience of auditory feedback. Based on previous studies of contingent feedback on movements in infancy (Lee and Newell 2013; Rochat and Striano 1999; Watanabe et al. 2011; von Hofsten 2004), these movement changes are assumed to reflect increased goal-directed behavior in anticipation of the auditory feedback, rather than simply an increase in spontaneous movements associated with increased arousal (e.g. fussy or crying; Shinya et al. 2019). This interpretation is also supported by our findings of the physiological measures.

For global changes of cardiac measures, the infants did not show any significant changes in heart rate and parasympathetic/sympathetic activity (i.e. HF, LF/HF of heart rate variability [HRV]) from Pre phase to Test phase. These results are consistent with the observation of a relatively stable heart rate during the Test phase (Fig. 2) and the previous study (Sullivan 2016). On the other hand, the HRV power spectrum should be interpreted cautiously considering recent discussions that the LF and HF components are not easily separated (e.g. Reyes del Paso et al. 2013). Given the limitation of these global measures, to reveal a more detailed physiological adaptation in the cardiovascular system, we further examined the event-related heart rate change (ERHR) regarding the limb movement inducing the onset of auditory feedback. As a result, through the experiment, infants inhibited the ERHR both before and after the onset of the auditory feedback, leading to the suppressed peak intensity of the ERHR.

Notably, the ERHR was found to be progressively suppressed even prior to the onset of the auditory feedback. Initially, during Pre phase, infants significantly increased heart rate before the timing of each limb’s acceleration exceeding the threshold (−1.7 sec). Previous studies in animals and human adults have reported that heart rate and arterial blood pressure increase immediately before or at the start of voluntary exercise (e.g. Komine et al. 2008; Ishii et al. 2016). Heart rate during voluntary motor exercise is controlled by cardiac autonomic activity resulting from central signals from higher brain centres (i.e. central command), although these regulations are also modulated by peripheral reflex arising from working skeletal muscle and arterial baroreflex (Matsukawa, 2011; Wiliamson et al. 2006). Therefore, the prior heart rate increase during Pre phase in this study is considered to originate from the central control of cardiovascular system associated with spontaneous limb movements. Then, during Test phase, the timing of ERHR occurrence was gradually closer to just before the onset of the auditory feedback, with a difference of more than one second between the Pre (−1.7 sec) and T4 (−0.5 sec) phases. In addition, the ERHR was significantly lower prior to the onset of the auditory feedback during T2-T4 phases than Pre phase. These findings suggest that the prior heart rate acceleration was suppressed in anticipation of the auditory feedback via central regulation of the cardiovascular system. Given that the changes in ERHR persisted beyond Test phase into Post phase (i.e. the period without auditory feedback), this persistent effect can be regarded as cardiovascular adaptation based on auditory-motor learning.

Our study does not provide direct evidence regarding what potential mechanisms have induced the suppression of ERHR prior to the auditory feedback via central regulation of the cardiovascular system. However, one possibility is that it is due to changes in attention based on the anticipation of auditory feedback. For the first two minutes of the Test phase, infants showed a rapid decrease in ERHR immediately after the auditory feedback (Fig. 3a) compared to Pre phase (Fig 3b). The heart rate deceleration may reflect orienting response and a level of sustained attention to the auditory stimuli (Lecanuet et al. 1992; Morrongiello and Clifton, 1984; Porges et al. 1973; Graham and Clifton 1966; Richards and Casey 1991). Additionally, the timing of the heart rate returning to the level of the surrogate data was slower gradually throughout Test phase (T1: 1.6 sec ∼ T4: 2.6sec). Nevertheless, the peak timing of ERHR was not changed throughout the whole experiment (i.e. about one second after the onset; Fig. 3a), indicating that the cardiovascular response itself to the limb movement is assumed not to have been delayed by the auditory feedback. Rather, considering that the suppression of heart rate increase before the onset of the auditory feedback was gradually strengthened after T1 phase, the timing of the heart rate deceleration can be considered to have been moved forward in anticipation of the auditory feedback. Heart rate deceleration accompanied by orienting response has been reported to occur not only when incoming stimuli are perceived, but also when they are anticipated or absent even in young infants (Colombo and Richman, 2002; Stamps, 1977). Therefore, it is possible that the gradual shift of the heart rate suppression from the post-onset to the pre-onset may reflect an earlier effect of the orienting response on heart rate due to increased anticipatory attention towards the auditory feedback.

Another possibility is that qualitative changes in the limb movements contributed to an absence of the pre-onset increase in the ERHR. As discussed above, our results of the significant changes in movement measures suggest increased goal-directed behavior in anticipation of the feedback based on previous studies (e.g. Watanabe et al. 2011). On the other hand, the occurrence of ERHR is thought to be associated with the limb movement related to the onset of the auditory feedback. Nevertheless, the movement may have been accompanied by some movements unrelated to the onset (e.g. other body movements, small limb movements under the threshold), especially during Pre phase and the early stages of auditory-motor experience. Therefore, increased efficient goal-directed behavior for the auditory feedback may be associated with inhibition of the movements unrelated to the auditory feedback, resulting in the absence of the heart rate increase prior to the onset. At this time, it cannot be ruled out that the anticipatory suppression of the ERHR may be due solely to changes in limb movements, since we evaluated only the timing for any limb’s acceleration exceeding the threshold regarding movement measures. Thus, future research should investigate the relations of more detailed movement measures to ERHR by using an accelerometer or a 3D motion-capture system.

In addition, we should consider the possibility that the averaging of overlapping ERHR may have caused attenuation in amplitude change and that the increase in frequency of limb movement over time may have produced ‘pseudo’ prior suppression and decrease in peak intensity of the ERHR. However, there were no significant differences in the frequency in the T1 to T3 phases as compared with the Pre phase (Fig. 2a). Thus, the prior suppression during T2 and T3 (Fig. 3b) phases and the decrease in peak intensity during T1 to T3 phases (Fig. 4) cannot be explained solely by the overlapping effect.

In taking a more comprehensive view of the above possibilities, the cardiovascular regulation in anticipation of the auditory feedback to limb movement can be interpreted in terms of allostasis. Allostasis, an extended concept of homeostasis, is a model for predictive regulation of the internal milieu through physiological and behavioral changes and requires anticipating physiological needs and preparing to satisfy them before they arise (Sterling 2012). The predictive regulation is achieved by the brain’s governing anticipatory behavior as well as by the regulating lower-level peripheral mechanisms based on prior knowledge. Here, it is of note that it may be difficult for young infants to anticipate auditory feedback associated with limb movements in a short period, since their ability to coordinate auditory and motor systems is still developing. Therefore, in this study, large prediction errors may have induced the rapid heart rate deceleration through orienting response (Hajcak et al. 2003), despite being at the peak of the heart rate response associated with the movement. Given these internal disturbances, the prior inhibition of the heart rate increase may contribute to dampening the rapid heart rate deceleration, resulting in reduced cardiovascular stress. In turn, the physiological needs may enhance auditory-motor learning and goal-directed behavior in young infants (Ecker and Gilead 2018; Nagai 2019; Verschure et al. 2014). Indeed, infants showed relatively stable heart rate and parasympathetic/sympathetic activity during Test phase, despite the increased frequency of the limb movement related to the auditory feedback. Therefore, it is possible that regulation of the cardiovascular system associated with attentional and movement changes may contribute to prolonged sound production and, in the long term, to acquiring more sophisticated musical behaviors.

Furthermore, our findings are also consistent with several pieces of evidence suggesting that music production and perception may involve arousal and stress regulation accompanied by changes in emotions and feelings (Bainbridge et al. 2020; Blood and Zatorre 2001; Habibi and Damasio 2014; Scherer and Zentner 2001; Chanda and Levitin 2013; Cirelli and Trehub 2020; de Witte et al. 2020; Tramo et al. 2011). Especially, it has been emphasized that auditory-motor synchronization plays a role in improving both the physiological load in cardiovascular system and exercise performance (Bood et al. 2013; Chaisurin et al. 2020; Trappe 2010). Reportedly, auditory-motor synchronization is an attention demanding process (Peper et al. 2012; Repp 2005). Thus, Bood et al. (2013) assumed that elevated attention on auditory-motor synchronization may contribute to distraction from fatigue and discomfort. This assumption is compatible with our interpretation of predictive regulation of cardiovascular system in young infants.

An additional limitation should be noted that the current study design did not include a control condition, although it is a pre-post design. Therefore, it is possible that the changes in ERHR may have been caused by increased arousal or fatigue over time, rather than the auditory feedback to the limb movements. On the other hand, we excluded infants who showed signs of excessive fussiness before Post phase to focus on the effects of the auditory back. In addition, we observed that infants did not show significantly higher heart rate during T1-T4 phases than Pre phase, although they did during Post phase. These facts suggest that the above possibility is relatively low, but still leave open the possibility that other factors over time may have influenced the changes of ERHR. Therefore, future studies should examine these possibilities, for example, by including a control condition without auditory feedback.

In conclusion, we revealed that 3-month-old infants regulate the cardiovascular system as well as limb movements in the early process of auditory-motor interaction. Through experiencing auditory feedback to the limb movements, infants inhibited the heart rate increase associated with the movements even before as well as after the onset of the feedback, leading to suppression of the peak intensity of heart rate increase. These findings suggest that emergence of auditory-motor interaction in young infants involves anticipatory regulation of the cardiovascular system accompanied by movement and attentional changes. The regulation may contribute to decreased stress on the cardiovascular system in auditory-motor interaction, providing a developmental basis of acquisition for more sophisticated goal-directed and musical behavior.

## Acknowledgements

We appreciate all the families for their participation in study. We also thank Kayo Sato, Yoshiko Koda, and Tomoko Yoneyama for their help with running the experiments. This work was supported by the Center for Early Childhood Development, Education, and Policy Research (Cedep), Graduate School of Education, the University of Tokyo; and the Grants-in-Aid for Scientific Research from the Japan Society for the Promotion of Science and the Ministry of Education Culture, Sports, Science and Technology (16H06525 to GT; 17KT0135 to HW; 20H04092 to SF; 20K14253 to YS)

## Statements and declarations

## Ethics approval

The study was approved by the ethical committee of Life Science Research Ethics and Safety, The University of Tokyo, and was conducted in accordance with standards specified in the 1964 Declaration of Helsinki.

## Consent to participate

Written informed consent was obtained from the participants’ parents.

## Availability of data and material

The data that support the findings of this study are openly available in Open Science Framework at https://osf.io/bc438/

